# Exact length distribution of filamentous structures assembled from a finite pool of subunits

**DOI:** 10.1101/042184

**Authors:** David Harbage, Jané Kondev

## Abstract

Self-assembling filamentous structures made of protein subunits are ubiquitous in cell biology. These structures are often highly dynamic, with subunits in a continuous state of flux, binding to and falling off of filaments. In spite of this constant turnover of their molecular parts, many cellular structures seem to maintain a well-defined size over time, which is often required for their proper functioning. One widely discussed mechanism of size regulation involves the cell maintaining a finite pool of protein subunits available for assembly. This finite pool mechanism can control the length of a single filament by having assembly proceed until the pool of free subunits is depleted to the point when assembly and disassembly are balanced. Still, this leaves open the question whether the same mechanism can provide size control for multiple filamentous structures that are assembled from a common pool of protein subunits, as is often the case in cells. We address this question by solving the steady-state master equation governing the stochastic assembly and disassembly of multi-filament structures made from a shared finite pool of subunits. We find that while the total number of subunits within a multi-filament structure is well defined, individual filaments within the structure have a wide, power-law distribution of lengths. We also compute the phase diagram for two multi-filament structures competing for the same pool of subunits and identify conditions for coexistence when both have a well-defined size. These predictions can be tested in cell experiments in which the size of the subunit pool or the number of filament nucleators is tuned.

## I. Introduction

Actin and tubulin are nanometer sized proteins that polymerize in cells to form filaments that are hundreds of nanometers to microns in length. These filaments often associate with other proteins to form larger structures that perform critical roles in cell division, cell motility, and intracellular transport.^1–3^ They are highly dynamic, namely they experience a high turnover rate of their constitutive protein subunits.^4,5^ Despite this, in order to function properly some of these structures (e.g., the mitotic spindle, or actin cables in budding yeast^5,6^) must maintain specific and well-defined sizes. Because these structures are assembled through a series of stochastic binding and unbinding events, the number of subunits in a structure is not uniform in time or within a population. Therefore, a structure with a well-defined size is one for which the distribution of the number of subunits it contains is sharply peaked at a non-zero value, or in other words, for which the fluctuations in their size over time, or over a cell population, are small compared to the mean size.

Many mechanisms for size regulation have been proposed based on careful experiments on cells and on reconstituted cellular structures *in vitro*.^6–11^ One such mechanism is to maintain a finite number of subunits within the compartment in which the structure is being assembled from the limited pool of subunits.^9^ Experimental evidence suggests that this mechanism is important in regulating the size of the mitotic spindle,^10^ as well as the size of nucleoli in a developing embryo.^12^ In this paper we develop a simple model of stochastic assembly and disassembly which we use to study this mechanism of size control in quantitative detail, with the ultimate goal of making testable predictions for future experiments.

The key assumption of the limited subunit pool model is the presence of a finite and constant in time number of subunits in a cell of constant size. In cells filaments do not assemble spontaneously in the cytoplasm but require specific proteins that act as nucleation sites for filament assembly; Figure 1(a). Within a single multifilament structure all the nucleation sites are equivalent, i.e., they all produce the same rate of filament assembly; Figure 1(c). An example is provided by actin cables in yeast which consist of many actin filaments bundled together all of which are nucleated and polymerized by the protein formin.^13^ We assume that the number of nucleation sites is constant within a single structure and that filaments only assemble at these sites; Figure 1(d). Note that this is a key difference with previously studied models of biopolymer assembly where it is assumed that subunits can freely associate with each other in solution to form filaments;^14^ this rarely, if ever, happens in cells.^15^

**Fig. 1:**
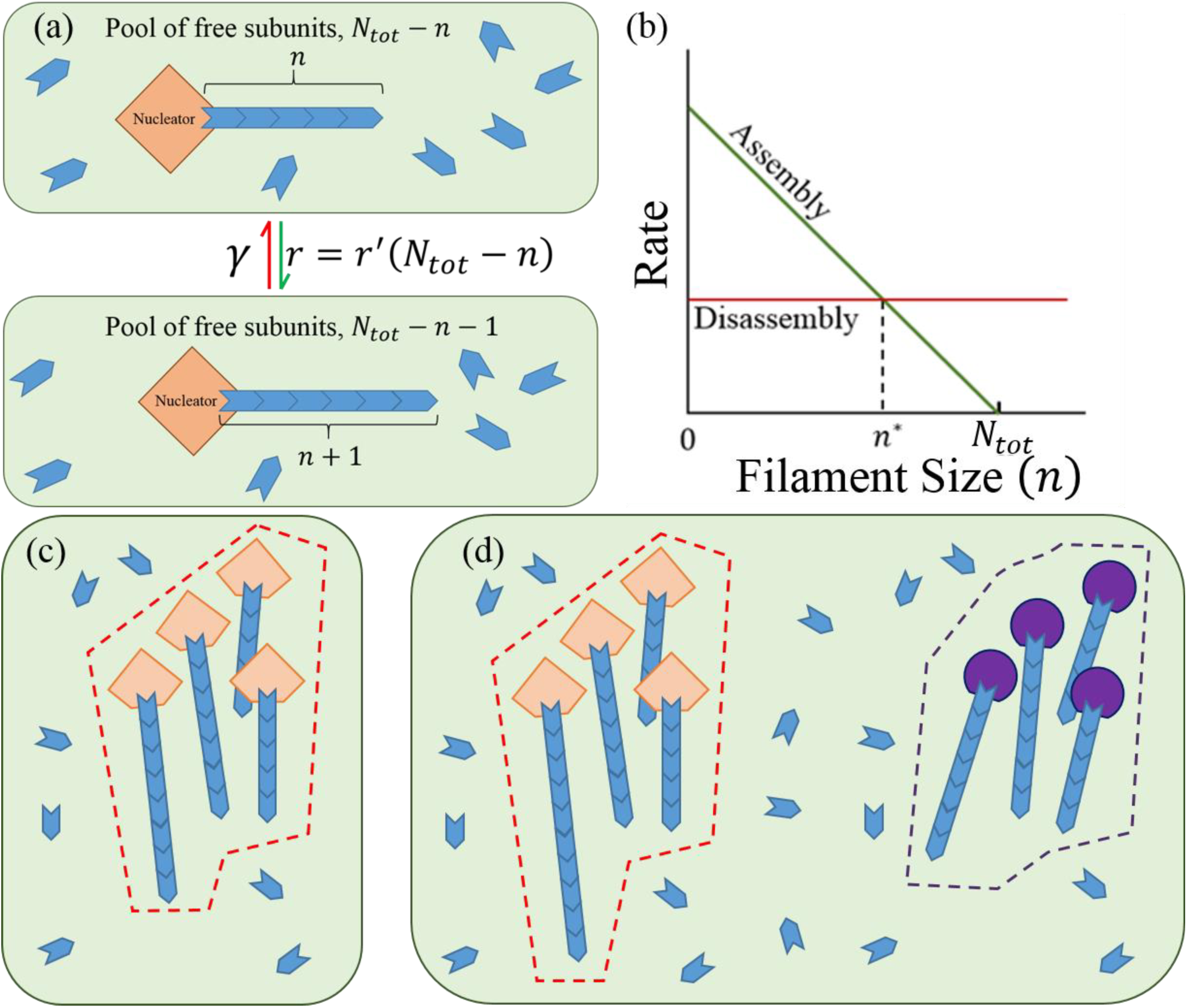
The limited subunit pool model. (a) A cartoon of filament assembly and disassembly for a single filament in a fixed volume (green box) containing a pool of *N_tot_* subunits. We assume that filaments are assembled only at the nucleator. (b) Rate of assembly and disassembly as a function of the filament length for a single filament being assembled from a pool of subunits whose total number is *N_tot_*; *n*^*^ is the filament length in steady state when assembly and disassembly match up. (c) One multifilament structure (bounded by the dashed red line). All of the nucleators in the structure are identical and have assembly dynamics described in part a. (d) Two multifilament structures in a common pool of subunits. Each structure is assembled by only one type of nucleators; the purple nucleators are characterized by different rate of assembly and disassembly than the orange ones.

The rate of subunit addition to a filament in a structure depends on the number of free subunits in the cell, or more generally, on their number in the compartment in which the structure is being assembled. As structures grow they deplete the cell of subunits and their growth rate slows. Eventually the growth rate matches the rate of subunit dissociation and a steady state with a constant number of subunits in filament form is reached; Figure 1(b). The stochastic nature of the assembly process leaves open the question: how large are fluctuations in size around the average? In other words, does the structure formed by the limited pool mechanism have a well-defined size?

The limited subunit pool model has been well characterized when there is only one filament assembling from a nucleation site. A closed form solution for the distribution describing the length of a single filament is easily computed using detailed balance. Depending on the values of the parameters that define the system it is possible to obtain length control of a single filament using this model. However, cytoskeleton structures in cells typically consist of many crosslinked and bundled filaments all made from the same protein subunit (e.g., actin or tubulin). Also, a single type of protein subunit may be used in the construction of a number of different structures depending on the regulatory proteins that associate with the filaments within the structure. Associated regulatory proteins include but are not limited to nucleators that act as sites for filaments to form (such as the Arp2/3 complex), elongators that increase the assembly rate of a filament (e.g., formins), cross-linking proteins that determine the topology of the structure (including *α*-actinin, filimin, and fimbrin), and degradation proteins that facilitate disassembly of filaments (cofilins).^16,17^ These observations motivate the key question we address herein, can a limited pool of subunits be used to regulate the size of multifilament structures?

## II. Methods and Theory

We consider a cell with *N_tot_* subunits and *N_f_* nucleators. Filaments can only assemble at the nucleators; Figure 1(c) and 1(d). Thus the filament associated with the *i*^th^ nucleator is simply referred to as the *i*^th^ filament. The *i*^th^ filament has a length *n_i_*, its disassembly rate is *γ_i_*, and, since we assume that the cell is a well mixed solution of subunits, the assembly rate is *r_i_* = 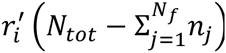 Namely, the assembly rate is proportional to the concentration of free subunits in the cell, which in a cell of constant volume is proportional to the number of free subunits 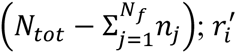 is the chemical second-order rate constant divided by the cell volume and it has units of sec^−1^. We also assume that the filaments do not interact in any way that would affect their rate of assembly or disassembly.

The key quantity of interest is *P*(*n_i_*; *t*) is the probability that the *i*^th^ filament has length *n_i_* at time *t*. Note that *P*(*n_i_*; *t*) is a marginal distribution of the joint distribution *P*({*n_i_*}; *t*). *P*({*n_i_*}; *t*) is the probability for the entire system to be in a state specified by the lengths of all the filaments, {*n_i_*} ≡ {*n*_1_,*n*_2_,…,*n_i_*,…,*n_N_f__*} at time *t*. The joint distribution *P*({*n_i_*}; *t*) is the solution to the master equation

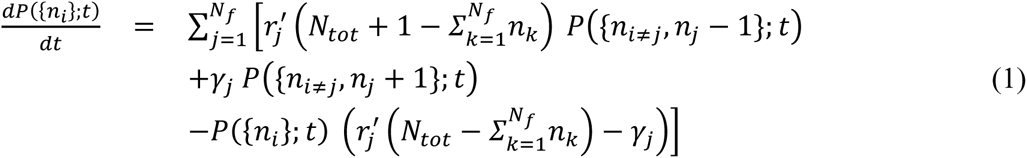

which simply enumerates all the changes in length that occur due to the elementary processes of subunit addition and removal from individual filaments, illustrated in Figure 1(a).

The rate of subunit turnover is typically fast compared to the lifetime of many structures,^18,19^ and we therefore solve Eq. 1 at steady state, when the probability distribution becomes stationary in time, i.e., when the time derivative in Eq. 1 is zero:

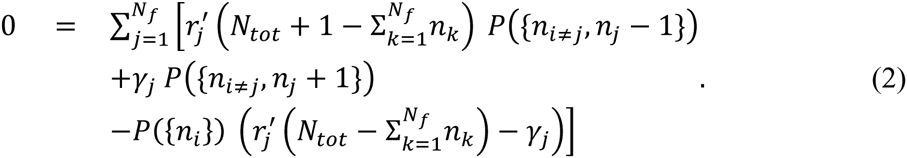

Here *P*({*n_i_*}) without the explicit dependence on *t* is the steady-state probability of finding the system in a state {*n_i_*}. Similarly, *P*(*n_i_*) will refer to the probability distribution of the *i*^th^ filament having length *n_i_* in steady state. In order to solve Eq. (2) for *P*({*n_i_*) it is helpful to rewrite it in the following manner:

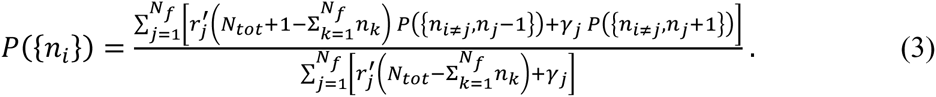

Eq. (3) is a system of linear equations. To solve for *P*({*n_i_*) we must also consider four additional constraints. First, if for any filament we have *n_j_* < 0 then *P*({*n_i≠j_*,*nj* < < 0}) = 0 i.e. filaments do not have negative length. Likewise, if the total number of subunits in all filaments is greater than the total number of subunits in the cell, 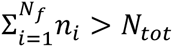, then 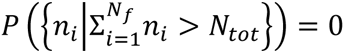 Third, we have normalization of the probability distribution,

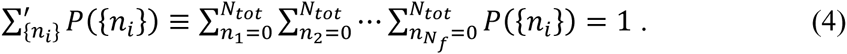

Finally, if *n_i_* = 0 then *γ_i_* = 0 because there are no subunits to remove from the filament. In all other cases is assumed to be a constant, independent of filament length. Eq. (3) maps to a well-known problem in queuing theory^20^ for which the solution is known to be in product form:

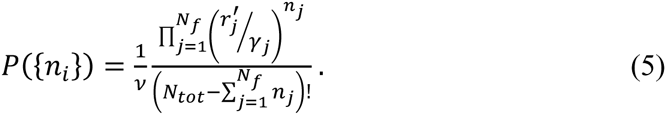

The quantity *v* can be computed from the normalization condition given in Eq. (4):

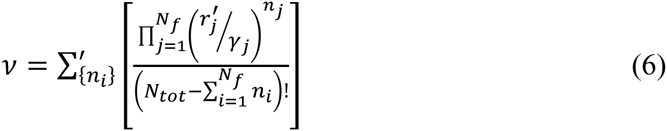

It’s a simple matter to check this solution by substituting Eq. (5) back into Eq. (3) and (4). Since Eq. (3) is linear in *P*({*n_i_*}) we conclude that the solution is unique.

We have already stated that cells employ networks of filaments for various tasks. Networks with different nucleators often have very different shapes that serve disparate roles in the cell.^1^ Because of this we are interested in collections of filaments assembled at identical nucleators. We call these collections multifilament structures; see Figure 1(c) where we illustrate two such structures each consisting of a number of filaments. The probability distribution for the sizes of structures can be constructed from Eq. (5). To differentiate structures from filaments we use Latin letter subscripts to denote filaments and Greek letter subscripts to denote structures. For each structure we define 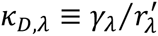 where 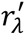 and *γ_λ_* are the assembly and disassembly rates for the filaments in *λ*^th^ structure, and *α_λ_* is the total number of nucleators/filaments associated with the *λ*^th^ structure. *κ_D_*,*_λ_* is the critical number of free subunits for a filament within the *λ*^th^ structure for which the rate of assembly and disassembly are equal. 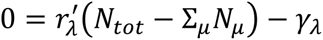 implies *κ_D,λ_* = *N_tot_* − Σ*_μ_N_μ_* = the number of free subunits. The critical number *κ_D,λ_* is also a dimensionless dissociation constant. It is related to the chemical dissociation constant *κ_D,λ_* (in molar units) for the chemical reaction of subunit addition to a filament shown in Figure 1(a), *κ_D,λ_* = *κ_D,λ_*/*V*_Cell_; here V_Cell_ is the volume of the cell (in liters). The total number of subunits in the *λ*^th^ structure is denoted *N_λ_* ≡ Σ*_j_n_j_* where *j* enumerates the filaments in the *λ*^th^ structure; *N_λ_* is the measure of the *λ*^th^ structure’s size that we adopt and the sought distribution of structure sizes is *P*({*N_λ_*}).

*P*({*N_λ_*}) is the sum of *P*({*n_i_*}) over all states {*n_i_*} that correspond to a state {*N_λ_*}. For each {*n_i_*} corresponding to a particular {*N_λ_*} we have the same number of free subunits, *N_tot_* − Σ*_i_n_i_* = *N_tot_* − Σ*_λ_N_λ_*. Also, while the values of each *n_i_* may vary between different {*n_i_*} we know that all *n_i_* corresponding to a particular *N_λ_* have *Σ_i_n_i_* = *N_λ_*. Thus we can rewrite Eq. (5) in terms of *N_λ_* and *κ_D,λ_*.

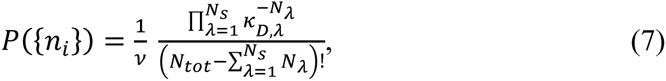

where *N_s_* is the number of structures in the cell (e.g., *N_s_* = 2 in Figure 1(d)).

From Eq. (7) we see that the sum *P*({*n_i_*}) over all {*n_i_*} corresponding to a given {*N_λ_*}, which gives the sought *P*({*N_λ_*}), has a constant summand. Hence to compute the sum we only need to count the number of states {*n_i_*} that correspond to a fixed {*N_λ_*} and then multiply that by the right hand side of Eq. (7). The solution to this counting problem is the binomial coefficient 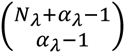 where *α_λ_* is the number of nucleators for the *λ*^th^ structure. Finally, we obtain for the distribution of multi-filament structure sizes in steady state

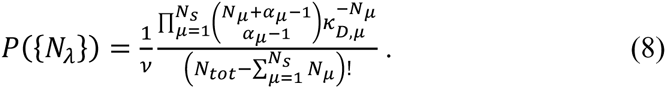

## III. Results

Eq. (5) and (8) are exact, closed-form steady-state probability distributions for the lengths of multiple filaments, and the sizes of multi-filament structures, all competing for the same finite pool of subunits. There are two special cases of Eq. (8) that are of particular interest given the kinds of structures that are found in cells. The first is a single multifilament structure (e.g., mitotic spindle), and second is two competing multifilament structures (e.g., actin patches and actin cables in yeast). We examine each of these cases below.

### Single Multifilament Structure

The single structure case is represented in our model by setting *N_s_* = 1 in Eq. 8:

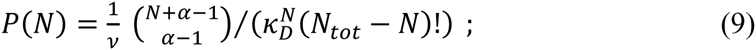

here *N* denotes the size of the structure, and *α* is the number of filaments/nucleators within this structure. This case also allows for a more concise formula of the normalization constant *v* in terms of special functions:

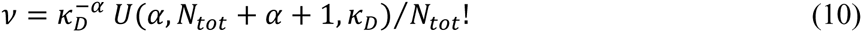

where *U*(*a*, *b*, *z*) is Tricomi’s confluent hypergeometric function.^21^

The marginal distribution *P*(*n_i_*) for the length of a single filament in this multi-filament structure comprised of *α* filaments is:

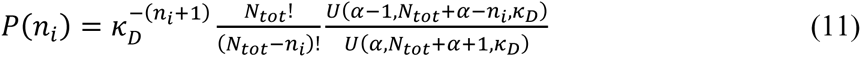

For *n_i_* < *N_tot_* − *κ_D_* there is a simple approximate form for this distribution *P*(*n_i_*)/*P*(0) ∝ ((*N_tot_* − *κ_D_* − *n_i_*)/(*N_tot_* − *κ_D_*))^*α*–2^, which we compare to the exact expression in Figure 2.

**Fig. 2:**
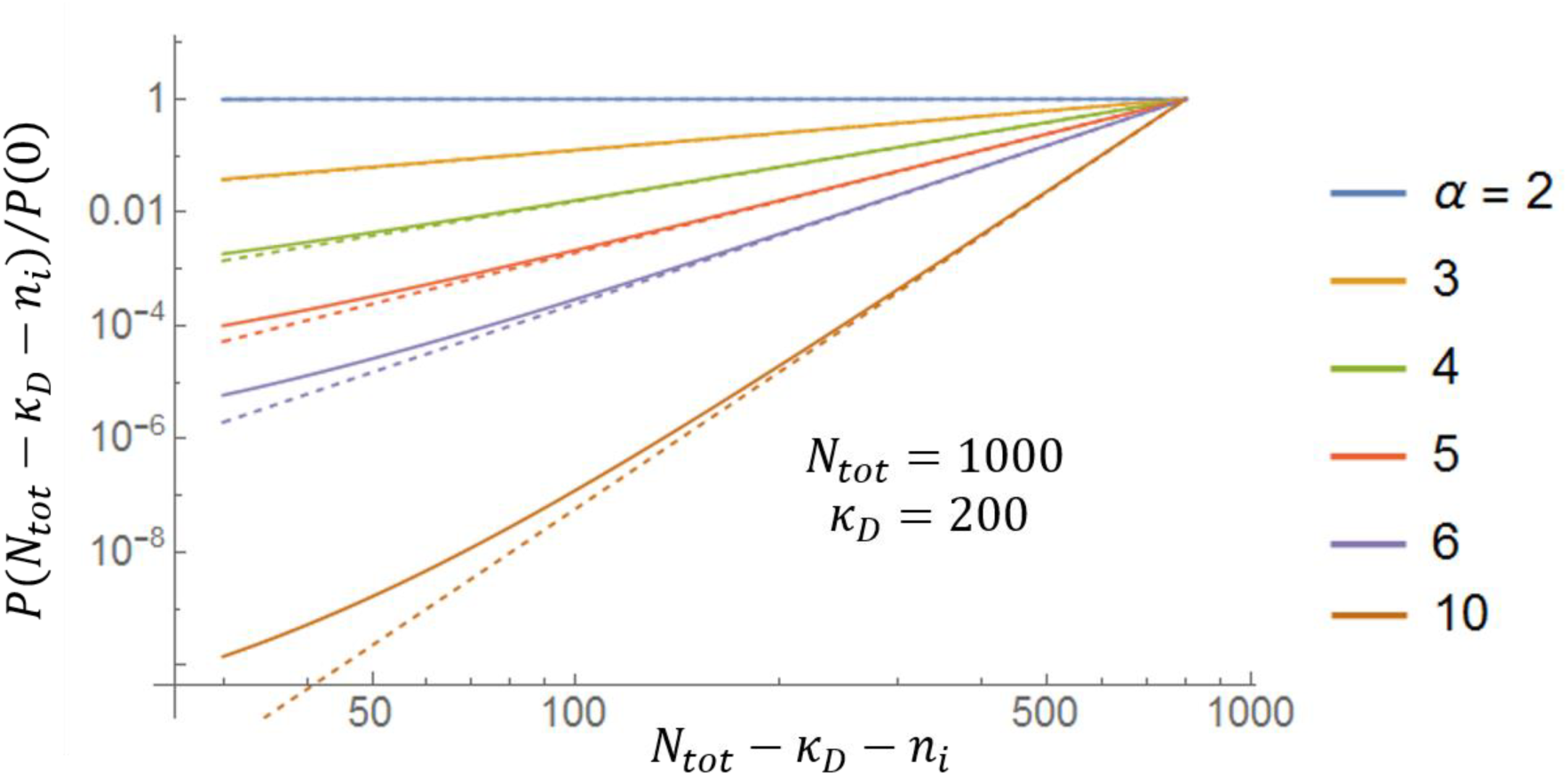
Exact and approximate (asymptotic) formulas for the length distribution of filaments in a multi-filament structure. Full lines correspond to the exact formula, Eq. (11), while the dashed lines are the asymptotic formula for a large number of subunits ((*N_tot_* − *κ_D_* − *n_i_*)/(*N_tot_* − *κ_D_*))^*α*–2^. Differences between the two formulas are not visible on a linear plot which is why a loglog plot is shown here.

We conclude that the filaments in a multifilament structure whose size is controlled by a finite pool of assembling subunits have a broad, power-law distribution of lengths. The power is given by the number of filaments in the structure (which is controlled by the number of nucleating centers) less two. A broad (fitting an exponential) distribution of microtubule lengths was recently measured for spindles assembled in frog extracts,^22^ in qualitative agreement with our findings. Clearly much more work needs to be done to make a more meaningful comparison with the theory presented here and experiments.

### Two Multifilament Structures

We examined the distribution of sizes for two structures each consisting of a number of filaments using Eq. (8) for a variety of model parameters. In Figure 3 we illustrate our results when we vary the number of filaments within a structure (*α_λ_*) and the dimensionless dissociation constant for the filaments within a structure (*κ_D,λ_*). For each of the two structures there are two possible outcomes depending on the values of the model parameters: either the structure has a well-defined size (illustrated in Figure 3 with a blue filament associated with a representative nucleator), meaning that the steady state distribution of the number of monomers within that structure is peaked around a non-zero value, or the distribution of sizes is monotonically decreasing with zero being the most likely size (illustrated in Figure 3 with a representative nucleator devoid of filament). Notably, we find regions of parameter space where the two structures, both with a well-defined size, can coexist. This shows that the finite subunit pool mechanism is capable of controlling the size of two multi-filament structures.

**Fig. 3:**
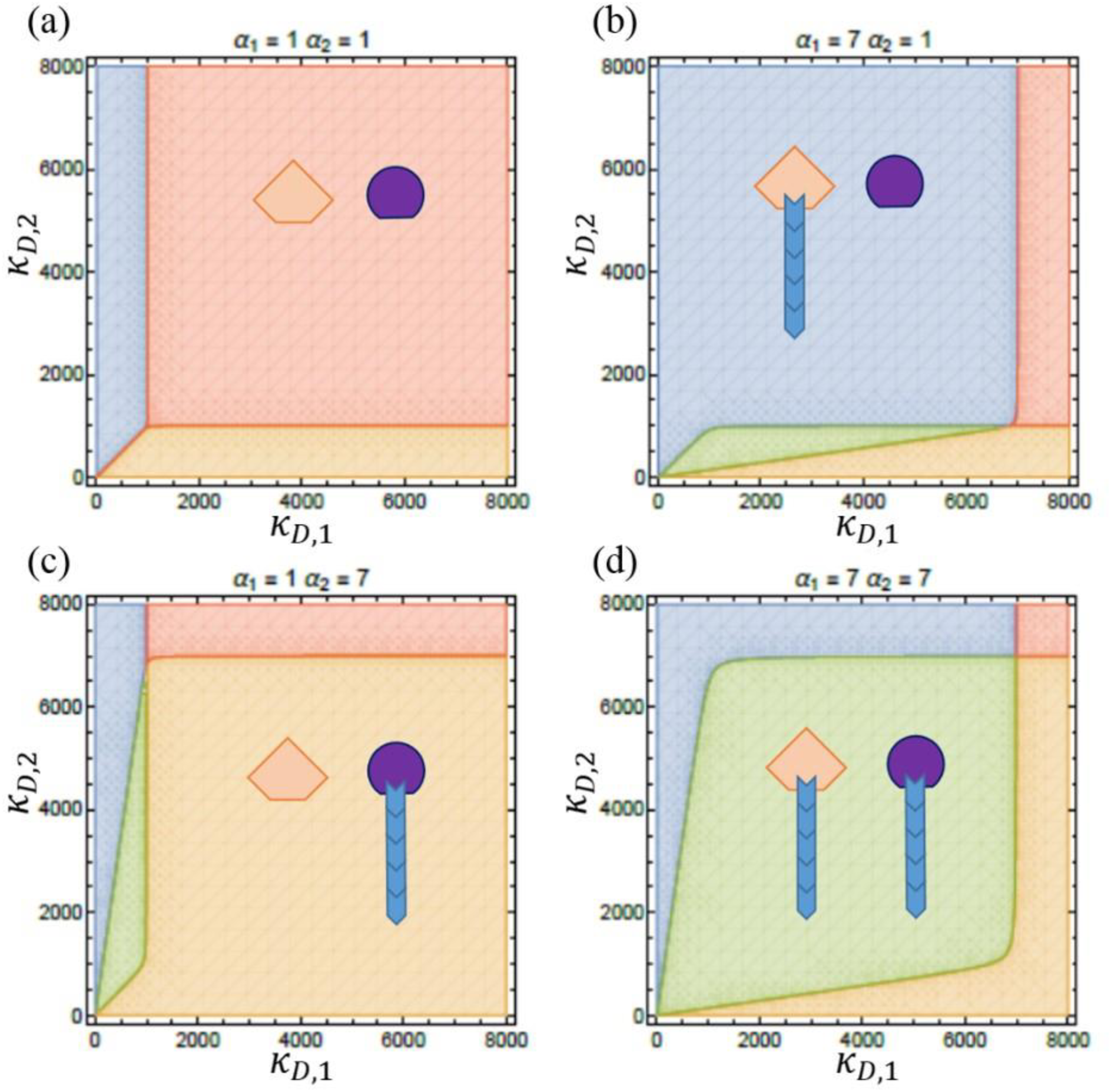
Phase diagrams for two multifilament structures. Blue regions correspond to parameter values for which the size of structure one is well defined, but not so for structure two. The orange region is when structure two has a well-defined size and structure one does not. In the red region neither structure has a well-defined size, and in the green region both structures have well-defined sizes. (a) For *α*_1_ = *α*_2_ = 1 when each of the two structures has only one filament there is no green region, i.e. the two filaments cannot both have a well-defined length in common pool of subunits. (b)-(d) As *α_λ_* is increased we see a region of coexistence (green) appear. We also see that the region where neither structure is well-defined (red) gets smaller. In all plots *N_tot_* = 1000 subunits.

Using the exact distributions described above we can also compute the dependence of the mean size of a structure on model parameters, as well as its standard deviation around the mean. We then characterize how well-defined a structure is by considering the noise of the size distribution of the structure, which is defined by the ratio of the standard deviation, 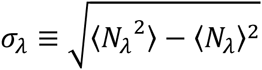, and the mean size 〈*N_λ_*〉. An exponential distribution is characterized by a noise of one, meaning that the fluctuations in size about the mean are equal to the mean size. For a structure to be well-defined its noise should be small compared to one.

In Figure 4 we show how mean structure size 〈*N_λ_*〉 changes with the total number of subunits *N_tot_* for a few different values of *α_λ_*, the number of filaments within a structure. In Figure 4(b) we observe that changing *N_tot_* changes the relative sizes of the two structures. For small *N_tot_* structure two is larger, but as *N_tot_* increases structure one becomes larger, i.e., it takes up more monomers from the subunit pool. Structure one with the lower critical number of subunits (*κ_D_*,_1_ < *κ_D_*,_2_) approaches a linear dependence of its size on *N_tot_* while the other structure plateaus to a value that’s independent of the total number of subunits.

**Fig. 4:**
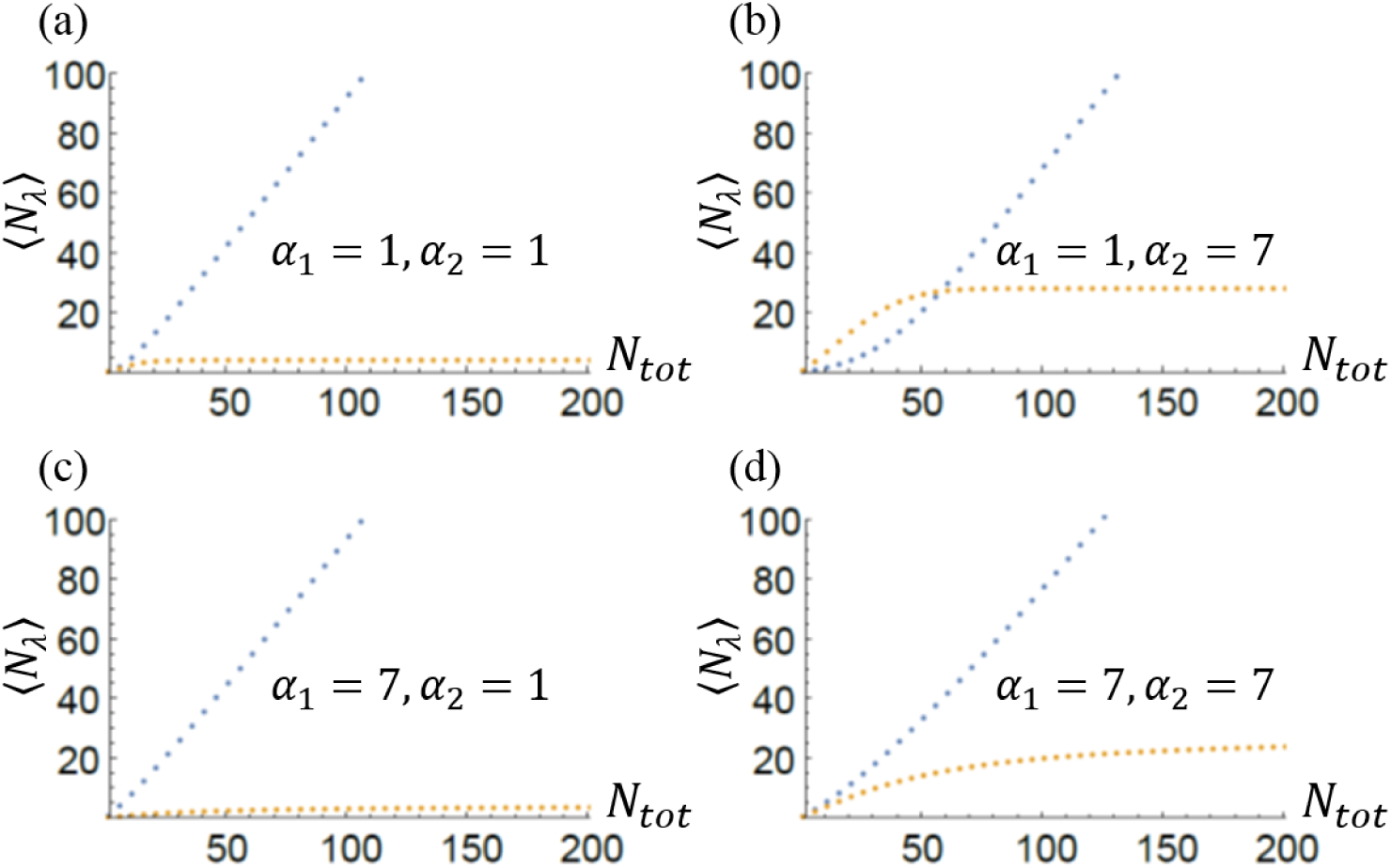
Mean sizes of two multi-filament structures competing for the same monomer pool. Blue line corresponds to structure one with *κ_D_*,_1_ = 4, and the orange line is associated with structure two with *κ_D_*,_2_ = 5. In all plots (a)-(d) we see that as *N_tot_* increases 〈*N*_1_〉 becomes linear in *N_tot_* while {*N_2_*) appears to approach a constant value. Comparing (a) and (c) with (b) and (d) we also conclude that increasing *α*_2_ increases the final value of 〈*N*_2_〉 as *N_tot_* becomes very large. It also appears that the increase in linear; for *N_tot_* = 200 in (a) 〈*N*_2_〉 = 4 while in (b) at *N_tot_* = 200 〈*N*_1_〉 = 28. This is a seven-fold increase in structure size as *α*_2_ goes from one to seven. This trend holds true for all values of *α*_2_ between one and seven.

Figure 5 shows the dependence of the noise on *N_tot_* for the same parameters as in Figure 4. In all plots the noise for structure one, which is the structure with the smaller critical subunit number, goes to zero as the total number of subunits increases while the noise for structure two goes to a non-zero constant whose value depends on the number of filaments in structure two (*α*_2_).

**Fig. 5:**
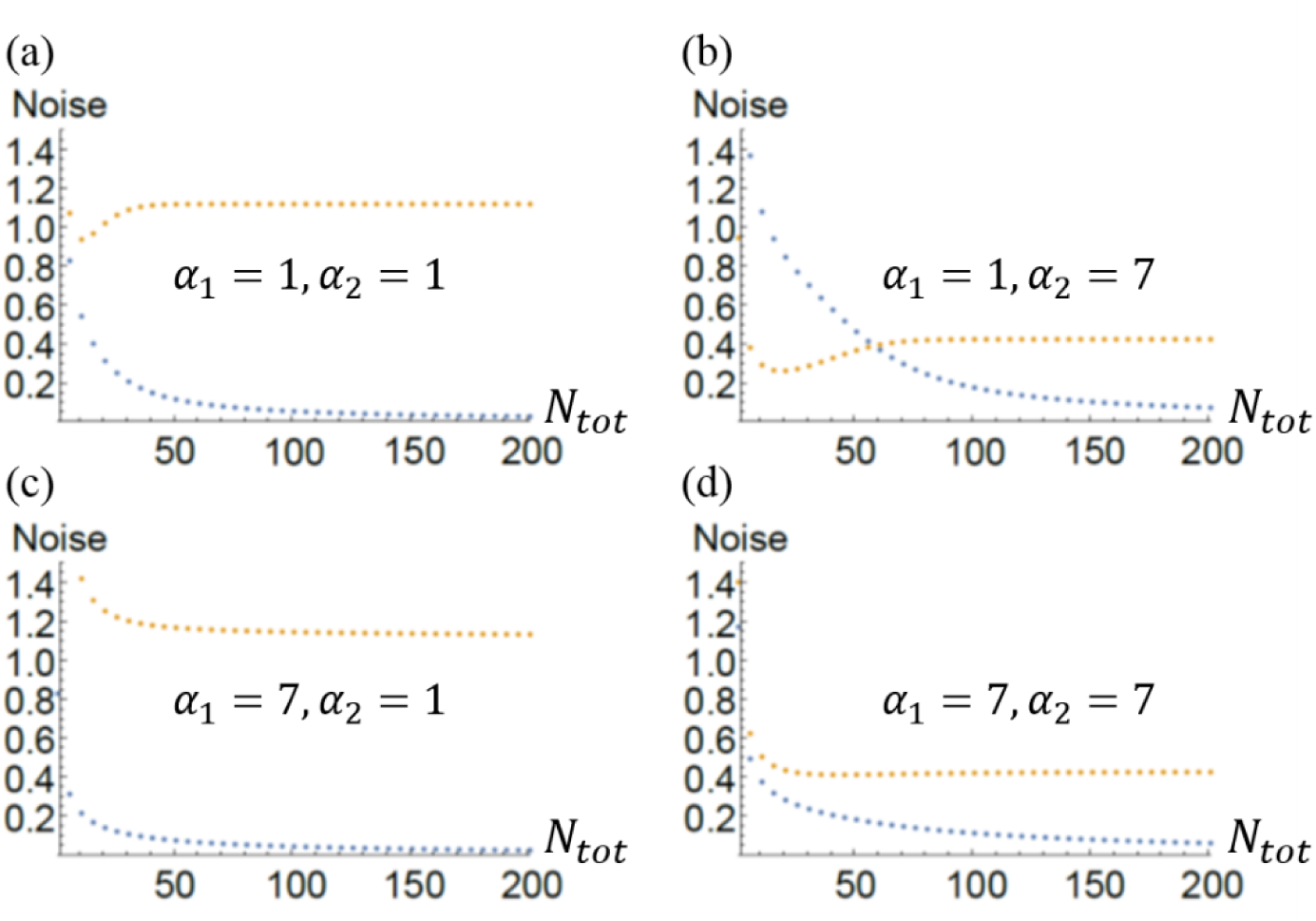
Noise of the size distribution for two multifilament structures. Blue is structure one with *κ_D_*,_1_ = 4 and orange is structure two with *κ_D_*,_2_ = 5. In (a) and (c) the noise for structure two is large (of order unity). This is expected since for *α*_2_ = 1 we expect structure two to be ill-defined since *κ_D_*,_1_ < *κ_D_*,_2_. In (b) and (d) the asymptotic value of the noise in structure two is less than one. In plots (a), (b), and (d) the noise for structure two drops to a minimum before increasing to its asymptotic value. In plot (c) it appears as though the noise for structure two drops monotonically to its asymptotic value.

## IV. Discussion

We have found exact closed-form solutions for the steady-state probability distribution of filament lengths in a system where growth is only limited by a finite pool of common subunits. From this distribution we found that a finite pool of subunits is not able to regulate the lengths of each individual filament. In spite of this, when multiple filaments are considered as parts of a multi-filament structure the overall size of the structure (i.e. the total length of all the filaments within the structure) can be well-defined. This means that despite the model’s simplicity, the finite pool of subunits is capable of simultaneously regulating the sizes of multiple multi-filament structures. Also, because of this model’s simplicity it is possible to make testable predictions about filamentous protein structures *in vivo* and *in vitro*. In particular, we find that the total number of subunits, *N*_tot_, number of nucleators for a structure, *α*_λ_, and the ratio of disassembly rate to assembly rate (the critical number of subunits), *κ_D,λ_*, all play a role in the mean size and variability of structure sizes. Intriguingly, recent experiments on actin structures in yeast have also found that these parameters are relevant to the regulation of structure sizes.^16,17^

While the calculations presented in this work address filamentous structures the conclusions that we draw hold more generally for three-dimensional structures as long as subunits are limiting and the rate of their addition to a growing structure is governed by its free concentration in solution. For example, recent experiments have shown that the size of the mitotic spindle reconstituted in cell extracts^10^ and the size of nucleoli in a developing embryo^12^ are both controlled by a finite pool of subunits. Notably, it was demonstrated that making the cell larger in a situation when the number of subunits is constant leads to smaller nucleoli, exactly as predicted by the finite pool mechanism.^12^

The predictions of the finite subunit pool model described above could also be tested *in vitro*, for example, by making use of beads with formins (actin nucleators) attached to them. Then the number of nucleators *α_λ_* could be controlled by tuning the number of formins on the bead.^23^ The total number of actin subunits *N_tot_* could also be controlled by confining the beads to microfluidically defined droplets of different sizes that have a fixed concentration of actin. Filament lengths can then be measured once steady state is reached and compared to the predicted length distribution.

Testing the model *in vivo* is more difficult but we can imagine controlling the number of actin nucleators in a yeast cell by regulating the expression of the gene that codes for the nucleator protein, or by regulating other proteins that are required to activate the nucleator. Another idea might be to take advantage of the natural variation of size in a population of cells, or changes in size in the process of development.^12^ Assuming a constant concentration of subunits this would allow us to test predictions of the model based on changes to *N*_tot_. We hope that the theory presented here will inspire these and similar types of *in vitro* and *in vivo* experiments, which will further our understanding of size regulation of subcellular structures.

## Acknowledgements

We are grateful to Bruce Goode and to David Kovar and members of their labs for many useful conversations. This research was supported by the National Science Foundation through the Materials Research Science & Engineering Centers grant number DMR-1420382.

---

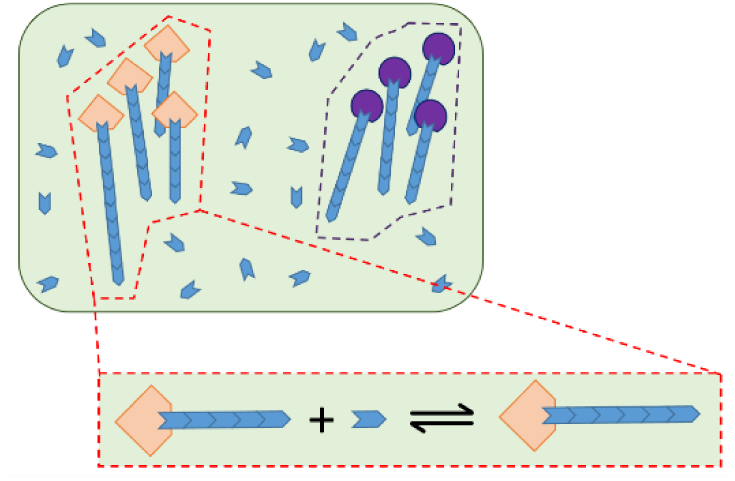
For Table of Contents Only

